# ALaSCA: a computational platform for quantifying the effect of proteins using Pearlian causal inference, with an example application in Alzheimer’s disease

**DOI:** 10.1101/2022.10.31.514546

**Authors:** Nina Truter, Zuné Jansen van Rensburg, Radouane Oudrhiri, David D. Van Niekerk, Ben Loos, Raminderpal Singh, Carla Louw

**Affiliations:** incubate.bio, London, United Kingdom; Department of Biochemistry, Faculty of Science, Stellenbosch University, Stellenbosch, South Africa; Department of Physiological Sciences, Faculty of Science, Stellenbosch University, Stellenbosch, South Africa

## Abstract

**Introduction:** An urgent need to delay the onset of aging-associated diseases has arisen due to increasing human lifespan. A dramatic surge in the number of identified potential molecular targets that could promote successful aging, has led to the challenge of prioritizing these targets for further research and drug development. In our previous work, we prioritized genes associated with aging processes based on their similarity to known aging-related genes and dysfunction marker genes in *C. elegans*. The goal of this study was to demonstrate the ability of our computational platform to identify molecular drivers of neuronal aging using specialized causal inference techniques. S6K was highly ranked in the previous study and here the nearby neighbors in its protein interaction network were selected to explore ALaSCA’s (Adaptable Large-Scale Causal Analysis) ability to identify possible drivers of Alzheimer’s disease.

**Methods:** Utilizing head and brain proteome data, two of ALaSCA’s capabilities were used to understand how protein changes over the lifespan of *Drosophila melanogaster* affect a feature of neuronal aging, namely climbing ability:

- Pearson correlation analysis was used to assess the relationship between the changes in abundance of specific proteins associated (through protein-protein interactions) with S6K and climbing ability.
- Pearlian causal inference, required to achieve formal causal analysis, was used to determine which pathway, associated with proteins linked to S6K, has the largest effect on climbing ability and therefore to what degree these specific proteins are driving neuronal aging.

**Results and discussion:** Based on the correlation results, the proteins associated with *fz*, a gene encoding for the fz family of receptors that are involved in Wnt signaling, display an increase in abundance as climbing ability declines over time. When viewed together with the *fz* proteins’ strong negative causal value, it seems that their increased abundance over the lifespan of *Drosophila* is an important driver of the observed decrease in climbing ability. Additionally, expression of the genes FZD1 and FZD7 (*fz* orthologs) is altered in the hippocampus early on in Alzheimer’s disease human samples and in an amyloid precursor protein mouse model.

**Conclusion:** We have demonstrated the potential of the ALaSCA platform to identify and provide evidence behind molecular mechanisms. This capability enables identification of possible drivers of Alzheimer’s disease - as the human orthologs of the proteins identified here, through its Pearlian causal inference capability, have been linked to Alzheimer’s disease progression.

## 1. Introduction

Global life expectancy has been increasing without a corresponding increase in health span and with greater risk for aging-associated diseases such as Alzheimer’s disease. An urgent need to delay the onset of aging-associated diseases has arisen and a dramatic increase in the number of potential molecular targets has led to the challenge of prioritizing targets that promote successful aging. In our previous work, we prioritized genes associated with aging processes based on their similarity to known aging-related genes and dysfunction marker genes in *C. elegans* (Truter *et al*., 2022). This previous work primarily focused on associative analyses of the gene expression profiles of known aging genes and genes previously unknown to be involved in aging (quantified by mRNA levels), supplemented with domain knowledge of molecular pathways and gene orthologs.

The goal of the current study was to demonstrate the ability of our computational platform to identify molecular drivers of neuronal aging in our Alzheimer’s disease project using specialized causal inference techniques. The application of our computational platform, ALaSCA (Adaptable Large-Scale Causal Analysis), includes supervised machine learning, correlation and Pearlian causal inference and associative analyses (Figure 1).

**Figure 1:**
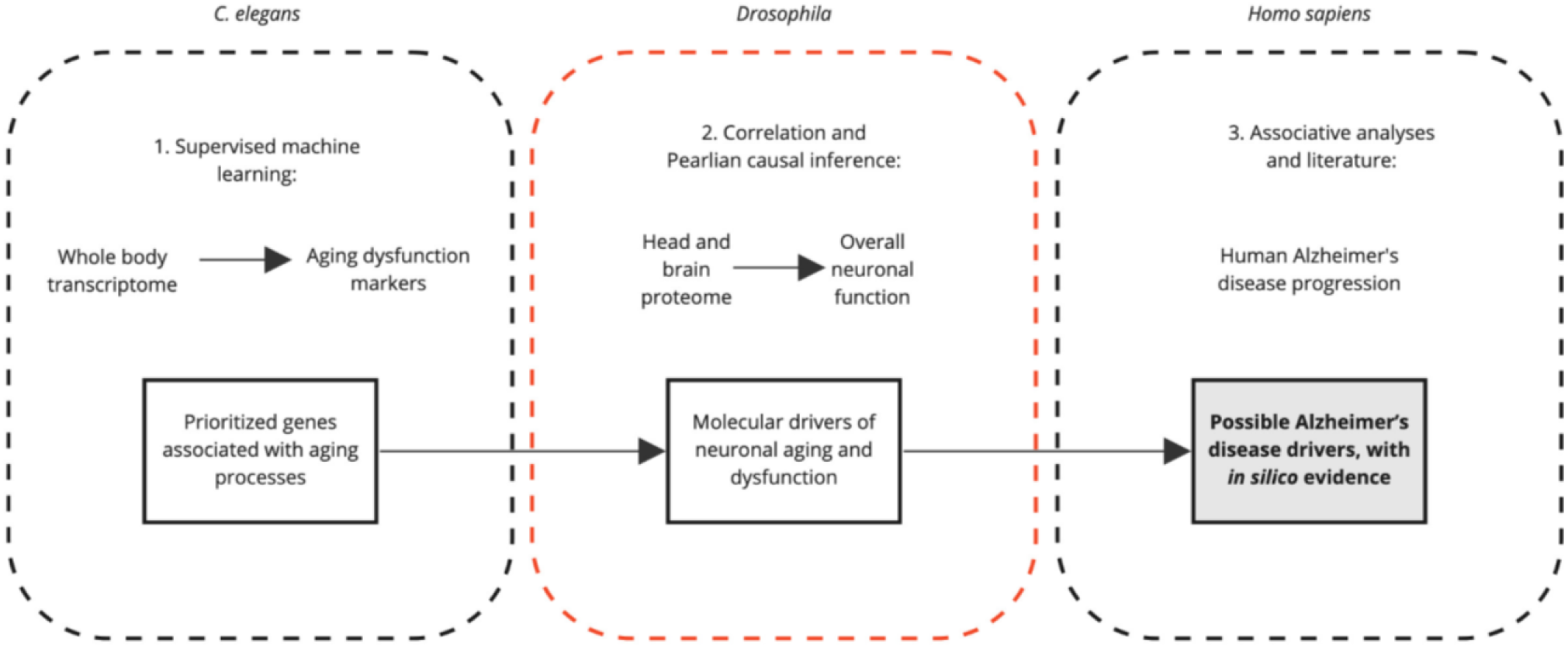
ALaSCA methodology to generate possible Alzheimer’s disease drivers. 1. Supervised machine learning was used to prioritize genes associated with aging processes through *C. elegans* transcriptome and dysfunction marker data (Truter *et al*., 2022). 2. In the current study (highlighted in red), we used these prioritized genes, correlation and Pearlian causal inference with *Drosophila* head and brain proteome data to identify the molecular drivers of neuronal aging. 3. The findings from these two system outputs are applied to Alzheimer’s disease progression data in *Homo sapiens* to generate possible disease drivers in humans.

In the current study, we investigated how the networks of proteins from the prioritized genes in our previous study could affect neuronal aging to provide further evidence for their potential value as targets for treatment to slow down Alzheimer’s disease progression, prior to *in vitro* and *in vivo testing*. The methodology includes understanding through which signaling pathway(s) these proteins drive neuronal aging, which could increase neuron vulnerability to certain insults that trigger neuronal dysfunction and Alzheimer’s disease risk in humans.

S6K was highly ranked in the previous study and here the nearby neighbors in its protein interaction network were selected to explore ALaSCA’s ability to identify possible drivers of Alzheimer’s disease. Although the previous approach enabled the prioritization of genes associated with aging, it did not describe how these genes and their associated protein networks could drive aging, specifically in the neuron. Given that S6K1 inhibition in mice reduced the generation of amyloid beta, a neurotoxic peptide which is found to aggregate in Alzheimer’s disease, our first aim was therefore to determine whether the S6K network has a causal effect on and therefore drives neuronal aging (Prüßing *et al*., 2013; Caccamo *et al*., 2015). Our second aim was to identify which signaling pathway with an association to S6K exerts the strongest effect on neuronal aging.

*Drosophila* serves as an improved model of neuronal aging with a more complex nervous system than *C. elegans*, which includes a brain consisting of 200,000 neurones that are biochemically similar to the human’s and functionally specialized substructures similar to the vertebrate central nervous system (Moloney *et al*., 2010; Prüßing *et al*., 2013; Barone & Bohmann, 2013). Molecular-level changes, such as that of protein abundances in neurons, are reflected in changes in climbing ability, an indicator of neuronal function (Ratliff *et al*., 2015). Climbing ability declines with age in a similar fashion to cognitive decline in the human (Barone & Bohmann, 2013).

Two of ALaSCA’s capabilities were used on these data to achieve our aims:

- Pearson correlation analysis was used to assess the relationship between the changes in neuronal abundance of specific proteins associated (through protein-protein interactions) with S6K and climbing ability.
- S6K is involved in many pathways that could affect climbing. Pearlian causal inference, required to achieve formal causal analysis, was used to determine which pathway, associated with proteins linked to S6K, has the largest effect on climbing ability and therefore to what degree these specific proteins are driving neuronal aging.

## 2. Methods

The Predicted Drosophila Interactome Resource was used to identify genes associated with S6K and subsequently their associated pathways that could affect climbing in *Drosophila* (Ding *et al*., 2020). Brain proteome data from Scholes *et al*. (2018) was merged with head proteome data from Sowell *et al*. (2007) and Brown *et al*. (2019) to obtain all quantified changes in abundance of proteins appearing in the S6K protein interaction network over 80 days. The datasets were scaled with Scikit-learn’s MinMaxScaler, whereafter a Gaussian Process Regression model was fitted for each of the proteins to infer any gaps in the data related to the sampling rate and ensure that the datasets align over the 80 days (Van Rossum *et al*. 2009 and Pedregosa *et al*. 2011). Climbing data were obtained from Long *et al*. (2014) and Ratliff *et al*. (2015).

Pearson correlations were used to determine the relationship between abundance changes of proteins from the S6K protein network and climbing ability. Next, we used causal inference methods described by Pearl (2009) and as we have previously demonstrated (Louw *et al*., 2022), to determine the effect of the S6K protein network on climbing over the lifespan of *Drosophila*. We then compared the effect of the S6K protein network to the effects of the protein networks of genes known to be related to the aging process, namely CG10702 (insulin signaling gene), Tor (mTOR) and aPI3K92E (PI3K) (Truter *et al*., 2022). These networks served as a benchmark of how strongly the proteins of known aging genes affect climbing. The absolute overall causal value was used here to compare the causal effects of proteins of aging genes on climbing. The combination of Pearlian causal inference with Pearson correlation analysis provides additional evidence to prioritize molecular mechanisms.

The positive and negative causal effects of the proteins associated with S6K were determined to identify which pathway most likely drives the decline in climbing ability. A positive causal value indicates an increase in the average height climbed in centimeters (and preservation of neuronal function) with an increase in protein abundance, whereas a decrease in this protein’s abundance would result in a reduction in the average height climbed. A negative causal value indicates a reduction in the average height climbed in centimeters (and decline in neuronal function) with an increase in protein abundance, whereas a decrease in this protein’s abundance would result in an increase in the average height climbed. The proteins with causal effect values larger than 0.3 or −0.3 were included for further analysis. Flybase was used to identify the human orthologs of genes associated with the proteins identified here (Gramates *et al*., 2022).

## 3. Results and discussion

### 3.1 The S6K network is a driver of neuronal aging

The S6K protein network displayed an overall effect on climbing (median = 0.208) that was comparable to that of the other networks from the three known aging genes, daf-2 (median = 0.228), let-363 (median = 0.228) and age-1 (median = 0.15) (Figure 2).

**Figure 2:**
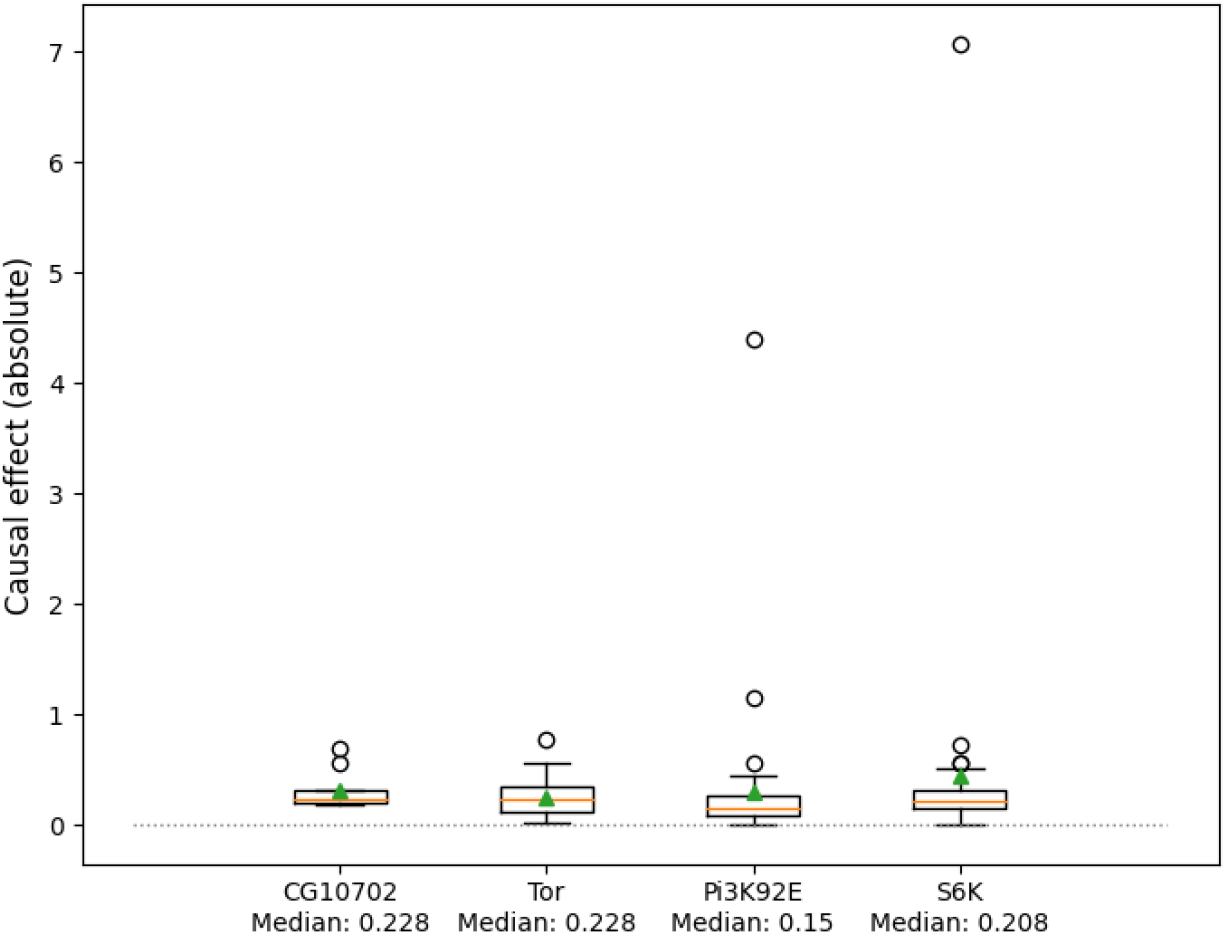
Box and whisker plots of the absolute causal effects of the protein-protein interaction networks linked to the four known aging genes on climbing ability in *Drosophila*.

### 3.2 Fz receptors of the Wnt signaling pathway have the largest causal effect on climbing

From the correlation analysis (Figure 3A), changes in abundance of the proteins encoded by *Gprk2*, *CG42366* and *PlexA* have the strongest positive correlation, and that of *fz* and *Ras85D* have the strongest negative correlation with the decline in climbing ability. However, it is unknown whether and how strongly changes in these proteins directly affect climbing ability or if these are spurious changes. Pearlian causal inference is able to provide strong evidence for each protein’s involvement in climbing (Figure 3B). Here, it is clear that the proteins encoded by *Gprk2* have a much larger positive causal effect on climbing ability than those encoded by *CG42366* and *PlexA*. Surprisingly, *fz*’s proteins have a dominant negative effect on climbing ability which overshadows the effect of proteins encoded by *Ras85D*. *Gprk2* and especially *fz* are therefore key to understanding how the S6K-network contributes to neuronal dysfunction, which leads to decline in climbing ability over the lifespan of *Drosophila*.

**Figure 3:**
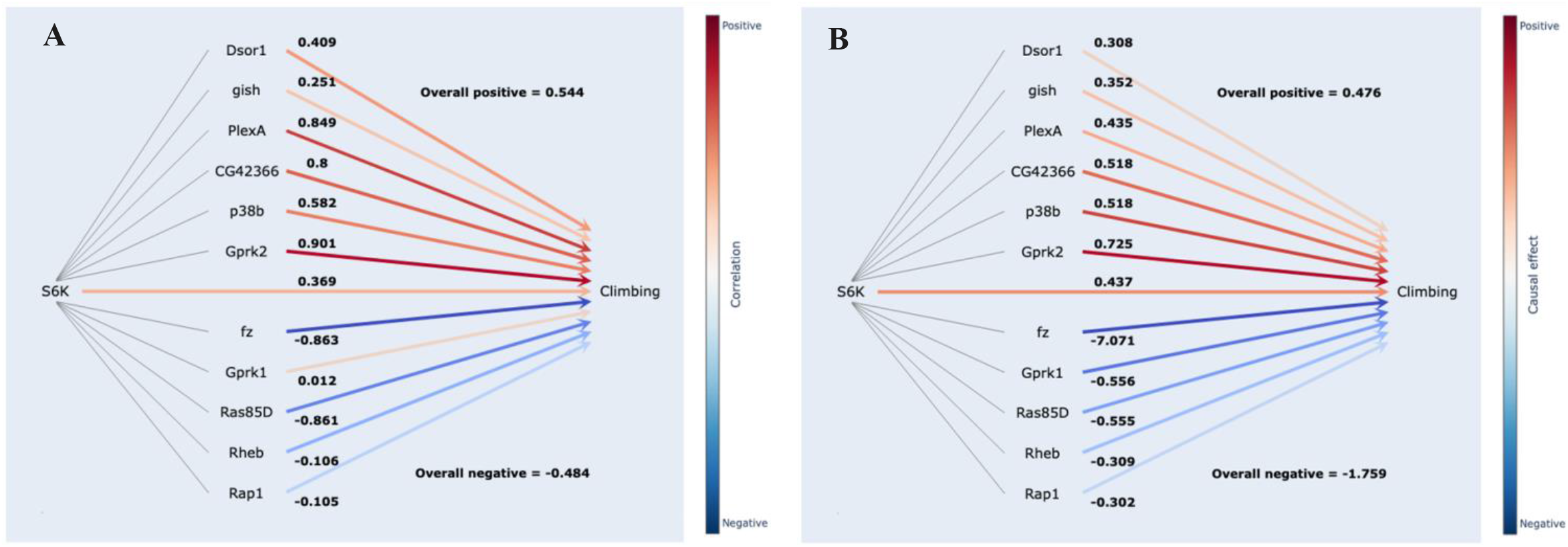
A) Correlation between abundance in S6K-associated proteins and decline in climbing ability. B) Causal effects of S6K-associated proteins on climbing ability.

By including a correlative approach, we can visualize the relative rate of protein abundance changes of these two genes compared to the decline in climbing ability, specifically around day 30. Just before day 30, the trajectory of each protein shows that the decrease in *Gprk2*’s protein abundance and increase in *fz*’s protein abundance corresponds to a very low climbing ability, which seems to improve as *fz*’s protein abundance continues to increase after day 30. Without this time-based correlative approach, the nuance in protein abundance changes before day 30 would be missed, as both *fz* and *Gprk2*’s protein abundance is the same (overlaps at day 28). How these protein changes could be driving the climbing ability decline is described by Pearlian causal inference. Although the strength of the correlation values between climbing and *Gprk2’s* (r = 0.901) and *fz*’s proteins (r = −0.863) (Figure 4), respectively, are similar (albeit in different directions), their causal scores (Gprk2 = 0.725 and fz = −7.071) show a 10X difference. This demonstrates the value of Pearlian causal inference in prioritizing which pathways are of most interest in understanding neuronal aging and deserves further study.

**Figure 4:**
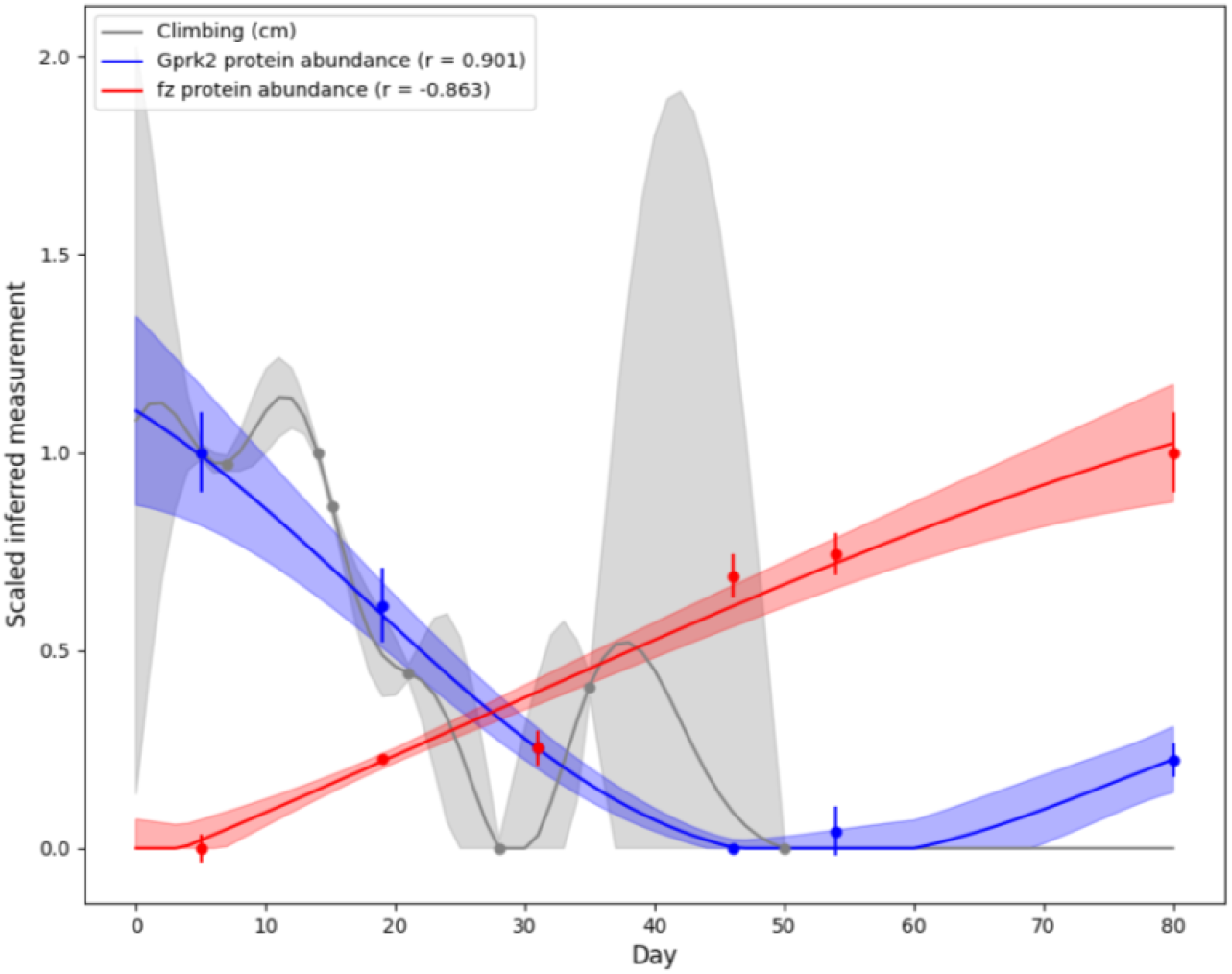
Protein abundance changes for Gprk2 (blue) and fz (red) over the lifespan of *Drosophila* (80 days) compared to climbing ability decline (grey).

*Gprk2* encodes for proteins that modulate G-protein coupled receptors and its elevated expression has been associated with Alzheimer’s disease in patients. The Gprk2 protein was found to promote Smo protein dimerisation and activity (Maier *et al*., 2014), and Smo proteins contain a fz class receptor, highlighting the involvement of fz-like receptors in understanding the decline in climbing. Because *fz* has a disproportionately large causal effect, we focus on providing a deeper understanding of its effects here. *fz* encodes for the fz family of receptors that are involved in Wnt signaling and specifically affects synapse formation in the central nervous system (Sahores & Salinas, 2011). Furthermore, Wnt signaling dysregulation is observed in both the aged human brain and Alzheimer’s disease patients (Palomer *et al*., 2019; García-Velázquez *et al*., 2017). Based on the correlation results, the proteins encoded by *fz* display an increase in protein abundance as climbing ability declines over time (Figure 4). When viewed together with the *fz* proteins’ strong negative causal value, it becomes clear that their increased abundance over the lifespan of *Drosophila* could be an important driver of the observed decrease in climbing ability. As Wnt signaling generally decreases with aging (Palomer *et al*., 2019), an increase in fz receptors may act as a compensatory mechanism to overcome this decline in signaling. However, fz receptors activate other pathways that affect Ca^2+^ signaling and cytoskeleton reorganization, which may contribute to neuronal dysfunction in the long term and could explain the reduction in overall neuronal function as seen in climbing ability (Wang *et al*., 2016).

*fz* was recently confirmed to be involved in the reduction of *Drosophila* climbing speed in aging flies (Gabrawy *et al*., 2022) due to its role in muscle development, maintenance, and aging. Here, knockdown of *fz* reduced control flies’ and drug-mediated rescue of climbing speed, indicating a contradictory positive effect of *fz* on climbing ability in this context. As we utilized head and brain proteome data, our findings, however, indicate that *fz* may play a role in changes in synapse formation and neuronal aging that ultimately drive a decline in climbing ability. Therefore, *fz* seems to affect climbing ability in a tissue-specific manner in *Drosophila*. Indeed, in early Alzheimer’s disease human samples and in an amyloid precursor protein mouse model, it has been shown that the expression of orthologs of *fz* (FZD1 and FZD7) in the hippocampus was altered (Palomer *et al*., 2022). The fz receptors were also implicated in the mechanism whereby soluble Aβ oligomers resulted in cognitive decline (Madhu *et al*., 2021).

The intimate involvement of *fz* in climbing and changes in its expression in Alzheimer’s disease, together with Gprk2’s increased expression in Alzheimer’s disease, helps to establish confidence in the capability of Pearlian causal inference combined with correlation to identify possible drivers of Alzheimer’s disease, as demonstrated here.

## 4. Conclusion

Through our computational platform we have previously identified proteins associated with aging-related processes that were associated with markers of dysfunction in aging. Here, we used correlation and Pearlian causal inference on a model of neuronal aging, to understand through which pathways these identified proteins could drive neuronal dysfunction that may lead to Alzheimer’s disease and to what extent these pathways could contribute.

We have demonstrated the potential of this platform (ALaSCA) to identify possible drivers of Alzheimer’s disease and their therapeutic potential. The human orthologs of both proteins identified here through Pearlian causal inference have been linked to Alzheimer’s disease progression. Because of the focus on aging processes in neuronal systems of simple organisms (*C. elegans* and *Drosophila)* and not merely protein changes as observed in Alzheimer’s disease *Drosophila* models, we could identify molecular drivers of neuronal aging. Upon subsequent mapping of these drivers to human orthologs, targets were identified that may improve the resilience of neurons to insults that could otherwise trigger neuronal dysfunction and Alzheimer’s disease risk in humans. This is enabled by the availability of transcriptomic, proteomic, and morphological time-based data over the lifespan of these organisms. Combining correlative and causal analyses enabled an unprecedented, in-depth understanding of protein changes in neurons, compared to either approach in isolation. Here, not only the changes of protein abundance in relation to climbing ability were described, but also to what degree these changes would affect climbing ability and therefore overall neuronal function. Pearlian causal inference therefore provides additional evidence to associative analyses on a protein or pathway’s involvement in neuronal aging, prior to its validation in *in vitro* or *in vivo* studies.

Future work for the Alzheimer’s disease program will include extending the analyses presented here, to more generally determine the proteins and pathways of aging processes that have the highest causal effect on different features representative of neuronal aging in *Drosophila*. These could be compared to the proteins and pathways that are altered in *Drosophila* Alzheimer’s disease models, to validate their contribution to pathology. These insights can be further related to human Alzheimer’s disease beyond comparison of orthologous genes by including associations with biomarkers derived from human brain data.

